# Attentional priority and limbic activity favor gains over losses

**DOI:** 10.1101/2024.08.12.607518

**Authors:** Kesong Hu, Eve De Rosa, Adam K Anderson

**Affiliations:** Department of Human Development, Cornell University, Ithaca, NY; Human Neuroscience Institute, Cornell University, Ithaca, NY; Department of Psychological science, University of Arkansas, Little Rock, AR

**Author notes:** **Correspondence:** To whom correspondence should be addressed to Kesong Hu or Adam Anderson.

**Keywords:** Loss aversion, Attention, Emotion, Regulatory focus, Mesolimbic, Reward, Ventral tegmental area, Insula

## Abstract

Prospect theory has suggested that decisions reflect a bias toward avoidance of loss compared to equivalent gains. In our study we examine whether a similar bias is found in decisions regarding whether a loss associated stimulus or a gain associated stimulus is given priority in perception. We find that for most people, gains are given priority over loss in the decision of which of the two stimuli occurs first. We also ask whether gains are reflected in greater activity in limbic systems related to emotion. In an fMRI study, we find that most people show greater emotional response to gains, not losses. We consider how these findings might be related to risk aversion in studies of decision making.

---

Studies of higher-level decisions have shown that “losses loom larger than gains”, meaning that the aversion to losing a certain magnitude is greater than the attraction to a gain of the same magnitude. For instance, in negotiations, eliminating losses is more effective than increasing gains(1), and most people reject gambles that offer a 50/50 chance of gaining or losing money, unless the amount that could be gained is at least twice the amount that could be lost(2). People are more sensitive to the possibility of losing objects or money than they are to the possibility of gaining the same objects or amounts of money (i.e., loss aversion). There is evidence that loss aversion is even present in monkeys, great apes, and children as young as age five(3–5). While loss aversion has received empirical support from a variety of studies(6–11), the source of loss aversion behavior and its variability remains poorly understood. Specifically, its role in directing attention with a higher priority to loss than to gain has not been tested. In the present study, we examine whether loss aversion bias exists in decisions regarding whether a loss-associated stimulus or a gain-associated stimulus is given priority in perception.

## Losses may draw more attention than gains

Often overlooked is that actual choices must be based on subjective perceptions of the attributes of the available options, and people may show distorted perception and rational inattention to gain or lose in choices(12). When facing a decision, people tend to choose the option which draws their attention, suggesting a causal role of attention in guiding loss aversion behavior. **Attention-based** theory predicts that losses draw more attention than gains (13, 14). For instance, loss attracts more automatic attention than gain (15), and loss aversion is reduced or eliminated where the biased allocation of attention to the loss does not occur(16). However, gain has also been shown to prioritize features for attention, dictate attention allocated to specific locations, and facilitate perceptual processing(17–20). So far, little is known about whether and to what extent losses are amplified relative to gains in directing attention. If the effect of gain and loss is rooted in basic human biases, it might be revealed in the most fundamental visual systems(21). If loss aversion reflects a more basic biological asymmetry in the importance of events to the organism, it should be reflected in the approach-avoidance/gain-loss system(22, 23).

## Losses may produce more emotion than gains

According to the **contrast-based** theory, loss aversion is rooted in an affective context, driven by increased weighting of losses compared to gains, and losses result in more extreme feelings(14, 24, 25). However, there are also findings that losses elicit less intensive emotions than gains (26–29). Emotion may reflect a fundamental feature of how gains and losses are assessed by the primate brain. Prior work suggests that loss aversion relates to activities in the amygdala, striatum, ventral anterior cingulate cortex (ACC), ventromedial prefrontal cortex (vmPFC), and the anterior insula(30–34). However, the current findings are mixed on whether the loss is associated with more activity than gains in the brain. For instance, insula is implicated in loss aversion(35), while it is inactivated to losses in an empirical study (10). Amygdala and ventral striatum are more activated for losses than gains in one study(36), while they are less activated or inactivated to losses than gains in the other study(10). It remains unknown whether loss produces more emotion and how the brain responds to gain and loss in directing attention.

Prior research on loss aversion is mostly carried out in high-level decision-making domains, including high-payoff game show decisions(37), financial markets(38), organ donation decisions (39), politics(40), trade policy(41), and intentional cognitive regulation(42), with the investigation of gain and loss processing taking place largely in an independent fashion. Moreover, previous findings are largely driven by the unequal probability weighting between gain and loss (43, 44). For instance, in gambling decisions, gaining and losing probabilities can vary, and typically the chances of winning, or of losing differ from 50%. *Unlike prior research*, in the present studies, we employ a dual-task paradigm, which includes a side judgment task and a temporal order judgment task, TOJ(45, 46). Gain and loss are equally reinforced in the side judgment task. ***The present study is unique*** because, rather than examining actual choice behavior (“experienced” utility) or anticipation of a positive or negative monetary outcome (“anticipated” utility) as prior studies have done, we investigate the sources of loss aversion, asking whether loss produces more attention, generates more emotion, and whether gain-loss contrast in perception reflects the engagement of distinct emotional processing in the brain.

Decision is not solely based on valuation but is dramatically susceptible to individuals’ preferences regulated by motivation systems(47, 48). Extending the basic hedonic principle, regulatory focus (RF) theory assumes that a person has two types of distinct motivational systems of goal pursuit, that is, the “prevention focus” regulates the behavior depending on the presence or absence of loss and the “promotion focus” regulates the behavior depending on the presence or absence of gain(48). We, therefore, further consider how **goal pursuit and value vary as a function of the RF orientation**, a promotion versus a prevention orientation, to understand how loss aversion arises,

## Results

### Study 1

Study 1 aims to assess the *attention-based theory* of loss. We start with one experiment (n=36), and then conduct a graphical meta-analysis with n=148 participants (87 females) across 6 experiments. The experiments are similar, and each consists of two intermixed trial types: *Side judgment* and *temporal order judgment* (TOJ) trials (**Fig.1**) (*SI Appendix, Supplementary Methods*).

**Figure 1.**
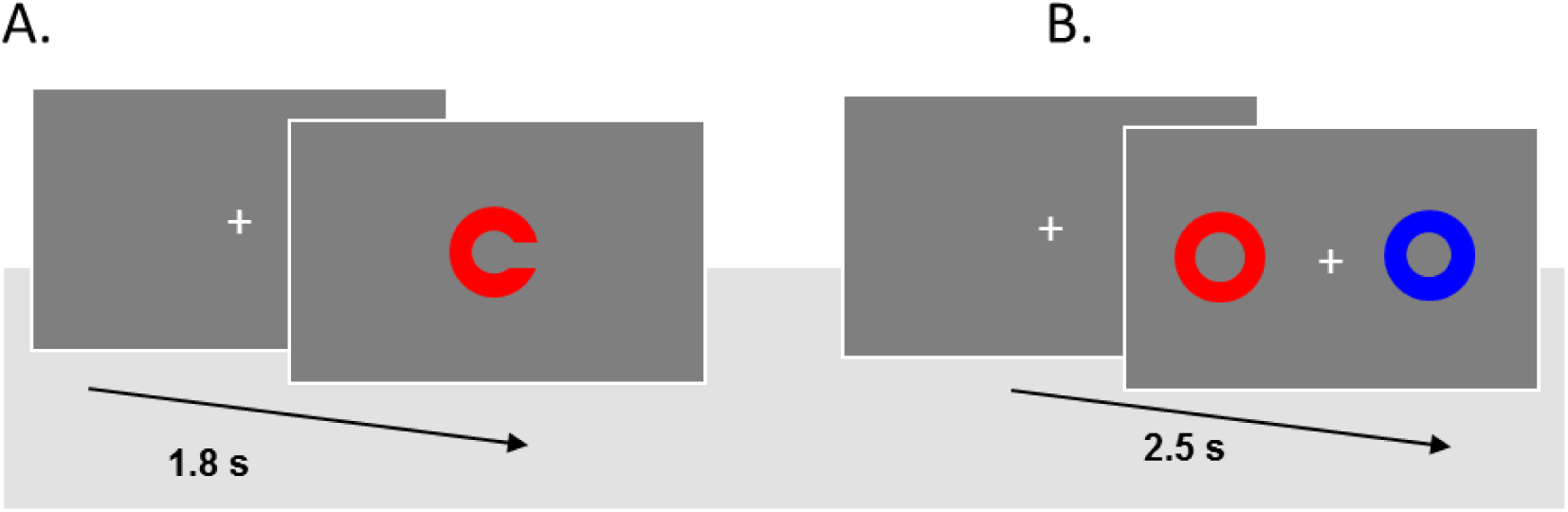
Example illustration of the trial sequences. The experiment consists of two tasks. (A) Side discrimination task. Each trial starts with an initial fixation display (1s), followed by a picture of a color circle image with a gap (0.8s). Participants respond to the side of the gap to gain or lose (avoid to lose) money. (B) Temporal order judgment (TOJ) task. Each trial starts with an initial fixation display (1s). Then a color circle (red, yellow, or blue) is presented either on the left or on the right side of the fixation, followed by a second different color circle, which appears on the opposite side of the first circle with a stimulus onset asynchrony, SOA (8, 18, 38, 68 and 98 msec). Participants indicate the temporal order of presented circles. Reinforcement and TOJ trials were pseudorandomly intermixed. See SI for details.

In the experiment with n=36 subjects, on average, participants have an equal proportion of gain (88%, $26.5) and loss (88%, $26.5), at the end of the experiment, t(35) =.25, p=.80, Cohen’s d=.04. Gain and loss are thus equally reinforced (**Fig.2A**). There is a significant main effect on RTs, F(2, 70) = 4.84, p=.01, ƞ^2^=.12, revealing that responses on gain and loss trials are faster than neutral. The ANOVA for error rates shows no significant main effect, F(2, 70) = 1.95, p=.15, ƞ^2^=.06(**Fig.2B**).

**Figure 2.**
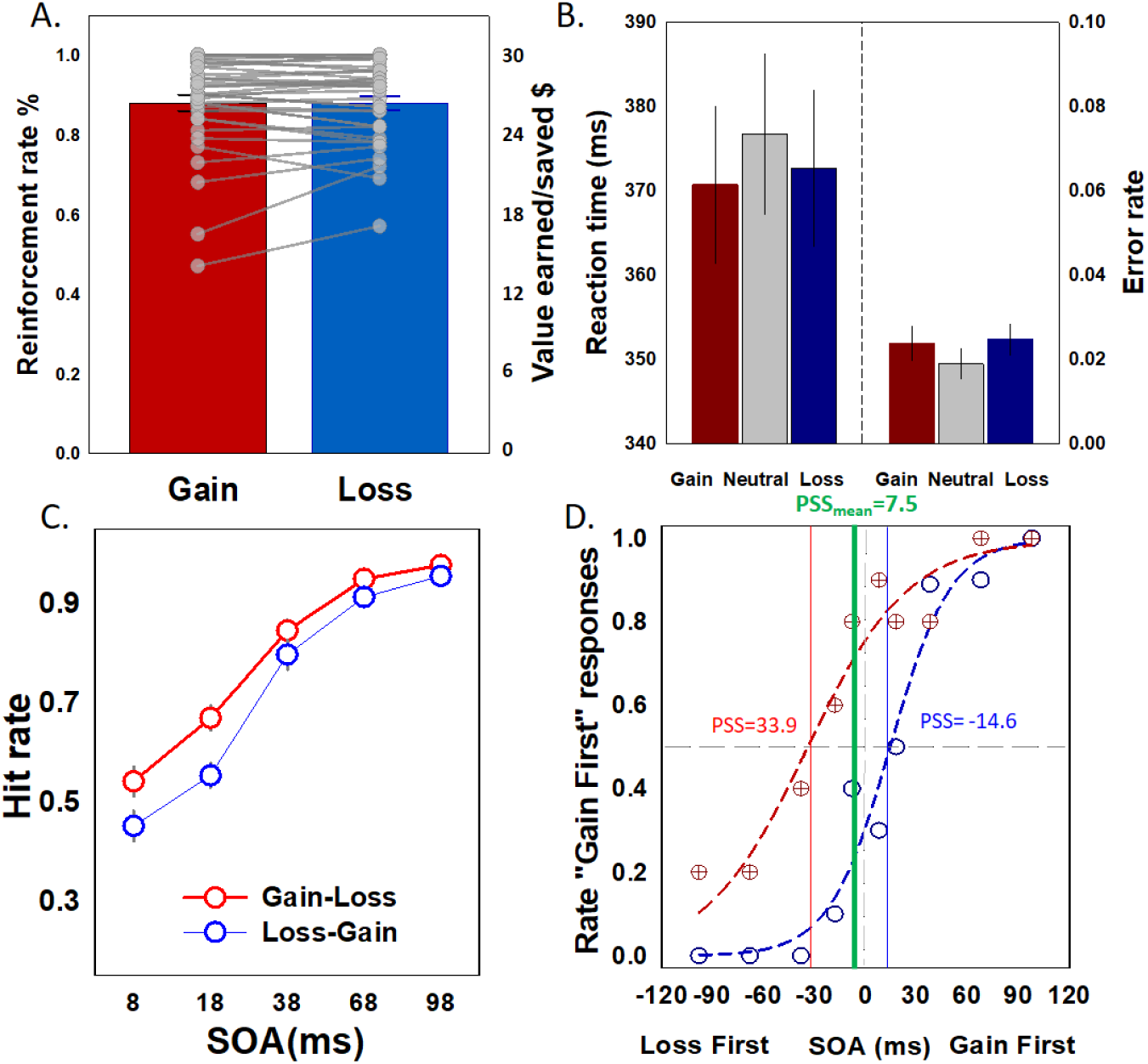
Reinforcement and temporal order judgment performance in the behavior experiment (n=36). A, Successful gaining (i.e., earned) and avoiding losses (i.e., saved) in rates and dollars. B, Mean RTs and error rates for gain, neutral, and loss trials in the value reinforcement task. C, the average hit rates for gain and loss from the “gain-first” and “loss-first” in gain-loss contrast TOJ task as a function of stimulus-onset asynchrony (SOA). D. Estimated point of subjective simultaneity (PSS) of all subjects (green), and for two subjects, one demonstrating a strong gain-first bias (red) and the other showing loss-first bias (blue). Error bars indicate standard errors of the mean (SEM).

To examine whether loss produces more attention than gain in perception, we first analyze accuracy, submitting hit rates to ANOVA [temporal order (Gain-first, Loss-first) x SOA (8,18,38,68, and 98 ms]]. A main effect of SOA, F(3,91) =200.27, p<.0001, ƞ^2^=.85, reveals that participants are much more accurate when the temporal difference between color circles is greater, and on average, for gain-first relative to loss-first colors, F(1,35)=11.44, p=.002, ƞ^2^=.25. The two –way interaction is not significant, F(3,105)=1.92, p=.130, ƞ^2^=.05(**Fig.2C**). ***Loss does not attract more attention***, which is further confirmed by the point of subjective simultaneity (PSS) estimation. We fit a Gaussian function to each individual’s data across onset asynchronies from the *TOJ task* (49). When gain and loss stimuli are pitted against each other, participants, on average, perceive loss as arriving 7.5 ms following gain, t(35) =3.12, p=.004, Cohen’s d=.52. However, the size is not large, and we observe substantial individual differences in the effect (**Fig.2D**). See *SI appendix* for the general description.

**Figure 3.**
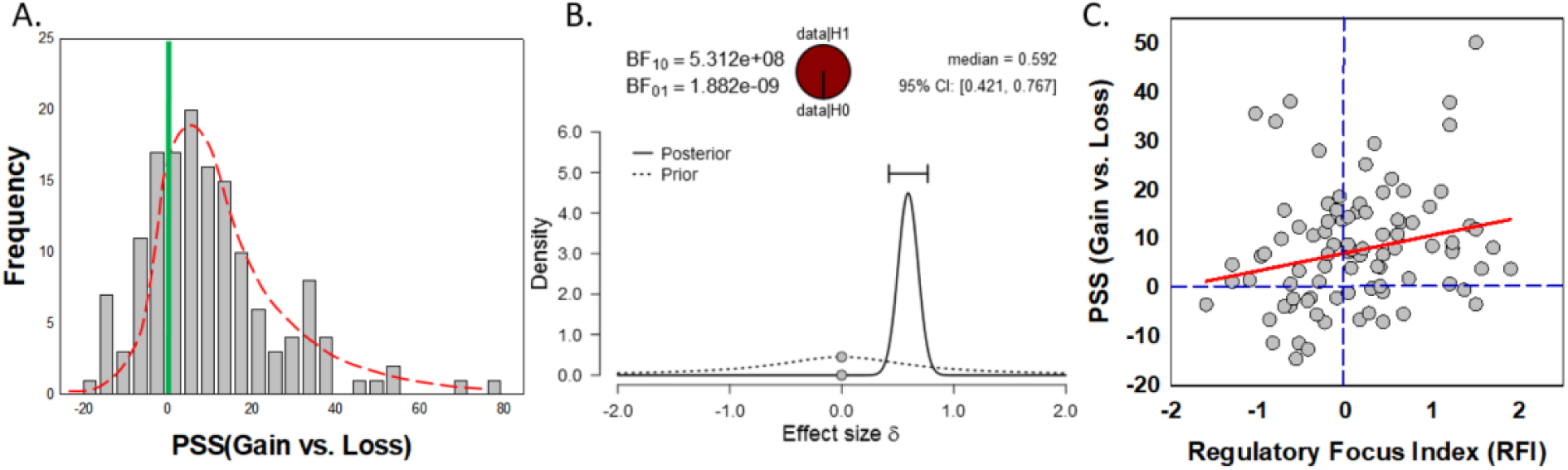
Distribution of individual differences in temporal perception of gain and loss cues (6 experiments, N=148 participants). A: Frequency distribution of temporal judgments for gain relative to loss cues (i.e., PSS_Gain vs.Lose_); B: Bayesian T-Test: Prior and posterior information. C. Individual differences in asymmetry in perceived temporal order were related to regulatory focus index (RFI= Promotion vs. Prevention). See text for details.

Small studies may be vulnerable to sampling variability, the random variation of an association across population subsamples(50). To specify the effect of loss aversion in perception, we next conduct **a graphical meta-analysis** with 6 experiments (**Fig.3A**). There is a skewed distribution in temporal perception away from zero towards gain color cues appearing first (Skewness,1.26; SEM: 0.20), but with a large variation (range, 100.4; Mean, 10.5; SEM, 1.41). To test deviation from what would be expected from chance, we contrast the distribution from a random null distribution based on resampling, using a permutation test(51) (permutation, 100,000): p<.00001, effect size, 1.20. This is further scrutinized via a comprehensive Bayesian analysis(52, 53). Bayes factor (BF_10_) represents a ratio between the likelihood of two hypothesizes (H_1_: PSS_Gain - Loss_>0; H_0_: PSS_Gain – Loss_=0 ) and how evidence supports one hypothesis over the other. When BF_10_ > 1, the greater its value, the stronger is the evidence for H_1_ when compared with H_0_. As seen in **Fig.3B**, there is greater evidence for H_1_ (BF_10_=5.31e +08), and a bias for preferential temporal perception of gain over loss cues. This bias occurs despite the fact that both gain and loss are equally reinforced, having a similar financial outcome in the monetary incentive task (permutation test, p=.443; BF_10_=.128) (**SI, Fig. 1**). Thus, similar performance in the value reinforcement task results in outsized attentional weighting of temporal perception of cues that signaled gain relative to loss. Although loss aversion is ubiquitous, **loss does not attract more attention than gain**.

There is, nevertheless, substantial individual variation in this gain-loss attentional bias (**Fig. 3A)**. We examine, within the framework of Regulatory Focus theory(47, 54, 55), whether this reflects an individual’s motivational orientation towards what is rewarding, differentially reinforcing the allocation of attention. The promotion versus prevention orientation, and their fit for different situations, lead to differences in the means of goal pursuit (54, 56). In the present study, the promotion and prevention orientations are recorded with 3 experiments (n=94 participants). To quantify anchor dependence, we first generate a differential RF index (RFI) (47, 55). Robust fit (to control for the influence of outliers) regression with PSS_Gain -Loss_ as the criterion variable, reveals individual differences in RFI are related to gain-loss asymmetry in perception of temporal order, Pearson’s r = .35, p=.001; robust R^2^ =.09, p =.005 (**Fig.3C)**. Individuals with strong promotion relative to prevention orientation demonstrate the most distorted temporal perceptions, with loss cues appearing to lag behind gain cues, i.e., loss posteriority, even when loss cues are presented first. By contrast, individuals with strong prevention relative to promotion orientation tend to show loss aversion. Differences in internalized **motivational orientation thus determine loss aversion**, shaping the differential salience of gaining and losing cues.

### Study 2

During multi-echo fMRI with the same paradigm as above, we test whether loss aversion has its roots in asymmetric emotional responses to losses than gains in perception. Study 2 shows similar behavior findings of gain-loss contrast, reported in full in ***SI Appendix***. Participants are equally reinforced with gain and loss, t(33)=1.51, p=.14, Cohen’s d=.22 (**Fig. 4A**). Examining participants’ temporal order judgments, we find that when gain and loss colors compete head-to-head, gain stimuli are judged to appear prior to the concurrently presented loss stimuli, t(33)=3.90, p<.001, Cohen’s d=.67 (**Fig. 4B**). In this subsample, RF orientation is associated with temporal perceptions (PSS_Gain vs. Loss_), Pearson’s r = .48, p=.007; robust R^2^ =.23, p =.009, with increased promotion relative prevention focus associated with reduced loss aversion (**SI, Fig. 3B**).

**Figure 4.**
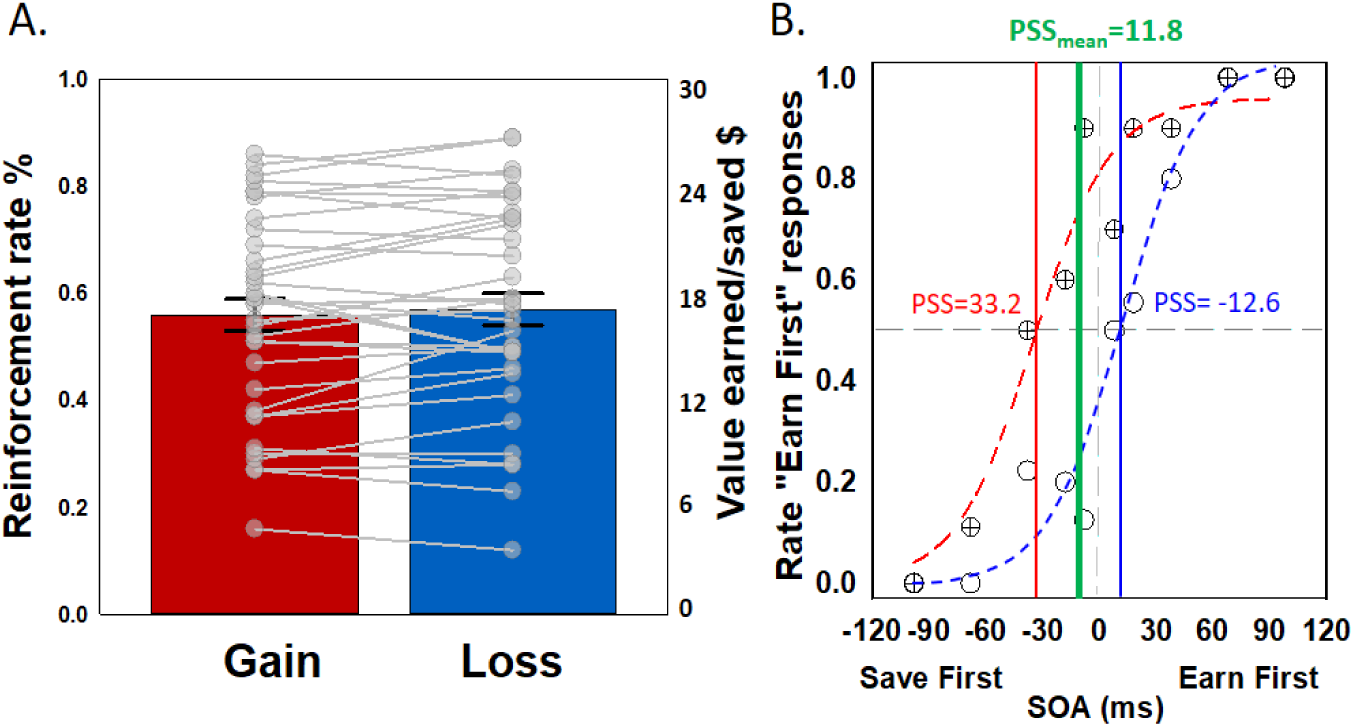
Reinforcement and temporal order judgment tasks performance during fMRI. A, Successful gaining (i.e., earned) and avoiding losses (i.e., saved) in rates and dollars in the value reinforcement task, and right panel, error rate; B. Estimated point of subjective simultaneity (PSS) of all subjects (green), and for two subjects, one demonstrating a strong gain-first bias (red) and the other showing loss-first bias (blue). Error bars indicate standard errors of the mean (SEM).

Consistent with a canonical reward response(57, 58), subcortical and cortical regions of the mesocorticolimbic system respond to *gain and lose* relative to neutral, including the ventral tegmental area (VTA), bilateral ventral caudate extending into nucleus accumbens (NAcc), bilateral anterior insula, medial orbitofrontal and medial prefrontal cortex (mPFC) (**Table 1**). These increases relative to neutral cues indicate that both gain and loss opportunities have positive utility and are rewarding and motivationally salient(59). The largely common reward system recruitment suggests that gain and loss opportunities reflect similar types of reward. **Loss does not produce more emotion than gain**. A direct contrast of loss and gain reveals, however, on average, a weak differentiation, with gain associated with greater recruitment in the putamen, thalamus, VTA, NAcc, caudate, and anterior cingulate cortex, while loss is associated with greater recruitment of the inferior frontal cortex (**Table 2**).

**Table 1.** Clusters that exhibited the effect of reward in voxelwise analysis at Whole-brain cluster-level corrected Alpha of .05 (Peak Talairach coordinates, t(33) values, and cluster size (k))

| Reward > Neutral |  |  |  |  |  |  |
| --- | --- | --- | --- | --- | --- | --- |
| Location |  | x | y | z | t | k |
| ***Occipital*** |  |  |  |  |  |  |
| lingual gyrus | L | -19 | -89 | -5 | 12.22 | 48 |
|  | R | 23 | -83 | -14 | 12.56 | 46 |
| ***Parietal/Temporal*** |  |  |  |  |  |  |
| Inferior parietal lobule | R | 44 | -50 | 51 | 24.05 | 198 |
| ***Frontal*** |  |  |  |  |  |  |
| Anterior insula | L | -28 | 18 | -2 | 12.19 | 133 |
|  | R | 46 | 15 | 1 | 21.51 | 621 |
| Medial Frontal gyrus | R | 5 | 15 | 45 | 15.33 | 279 |
| Middle Frontal gyrus | R | 41 | 45 | 18 | 18.11 | 621 |
| Anterior Cingulate cortex | R | 8 | 25 | 27 | 6.44 | 279 |
| ***Subcortical*** |  |  |  |  |  |  |
| NAcc/Caudate Nucleus | L | -7 | 3 | 9 | 8.85 | 77 |
|  | R | 8 | 6 | 7 | 13.74 | 331 |
| VTA | L/R | 3 | -18 | -12 | 5.69 | 331 |
| ***Cerebellum*** |  |  |  |  |  |  |
|  | L | -7 | -71 | -23 | 11.88 | 50 |
|  | L | -34 | -62 | -23 | 8.06 | 36 |

**Table 2.** Voxelwise analysis (Peak Talairach coordinates, t. (**33**) **values, and cluster size (k))**

| **Gain > Loss** |  |  |  |  |  |  |
| --- | --- | --- | --- | --- | --- | --- |
| **Location** |  | **x** | **y** | **z** | **t** | **k** |
| ***Frontal*** |  |  |  |  |  |  |
| Anterior Cingulate cortex | R | 5 | 42 | 9 | 4.04 | 10 |
| Middle Cingulate cortex | R | 5 | 36 | 27 | 5.67 | 4 |
| ***Subcortical*** |  |  |  |  |  |  |
| Putamen | L | -10 | 6 | -8 | 3.10 | 6 |
| Thalamus | R | 8 | -15 | 12 | 3.37 | 4 |
| VTA | L | -1 | -17 | -11 | 2.90 | 3 |
| NAcc | L | -13 | 9 | -8 | 2.67 | 3 |
| Caudate | R | 8 | 15 | 4 | 4.14 | 3 |
| ***Cerebellum*** | L | -31 | -59 | -23 | 5.52 | 8 |
| **Loss > Gain** |  |  |  |  |  |  |
| Inferior frontal gyrus | L | -43 | 42 | 9 | 9.75 | 12 |

P<.005 (uncorrected), k>=3 (81 mm<sup>3</sup>).

Despite their similarity, individual differences in gain and loss priorities may reflect the different magnitude of reward responses, rendering these opportunities differentially salient. When we analyze affective responses to the color cues pre- versus post-monetary reinforcement, individual differences, specifically in left insula response, predict differential subjective valence of gain relative to loss colors, Pearson’s r = .47, p=.008; robust R^2^ =.19, p =.013 (SI, **Fig. 4A**) and thus reflect differential subjective affective salience (60) of cues signaling to losses and gains.

Individual differences in VTA response predict the magnitude of distorted temporal perceptions of gain relative to loss cues. The greater the VTA response to gain relative to loss opportunities during monetary reinforcement, the greater the disparity in their temporal perception, PSS(Gain-Loss), Pearson’s r = .51, p=.003; robust R^2^ = .25, p =.004(**Fig. 5A**). In addition to the VTA (**Table 2**), the bilateral anterior insula, left, Pearson’s r = .54, p=.001; robust R^2^ = .28, p =.002 **(Fig. 5B)**, and right, Pearson’s r = .40, p=.027; robust R^2^ = .09, p =.105(SI, **Fig. 5**), predict which individuals demonstrate decreased salience of loss in temporal order judgments. Further, an individual’s VTA tuning to gain vs. loss is related to anterior insular response, Pearson’s r = .54, p=.001; robust R^2^ = .28, p=.002(SI, **Fig. 4B**). Note this relation is confirmed by trial-by-trial functional connectivity analysis(61, 62). The insula is a critical hub orchestrating salience in both internal feelings and external behavior and attention(63). In conjunction, greater VTA and anterior insular responses favoring gaining are associated with an individual’s degree of temporal distortion, supporting a relatively slowed perception of losing cues.

**Figure 5.**
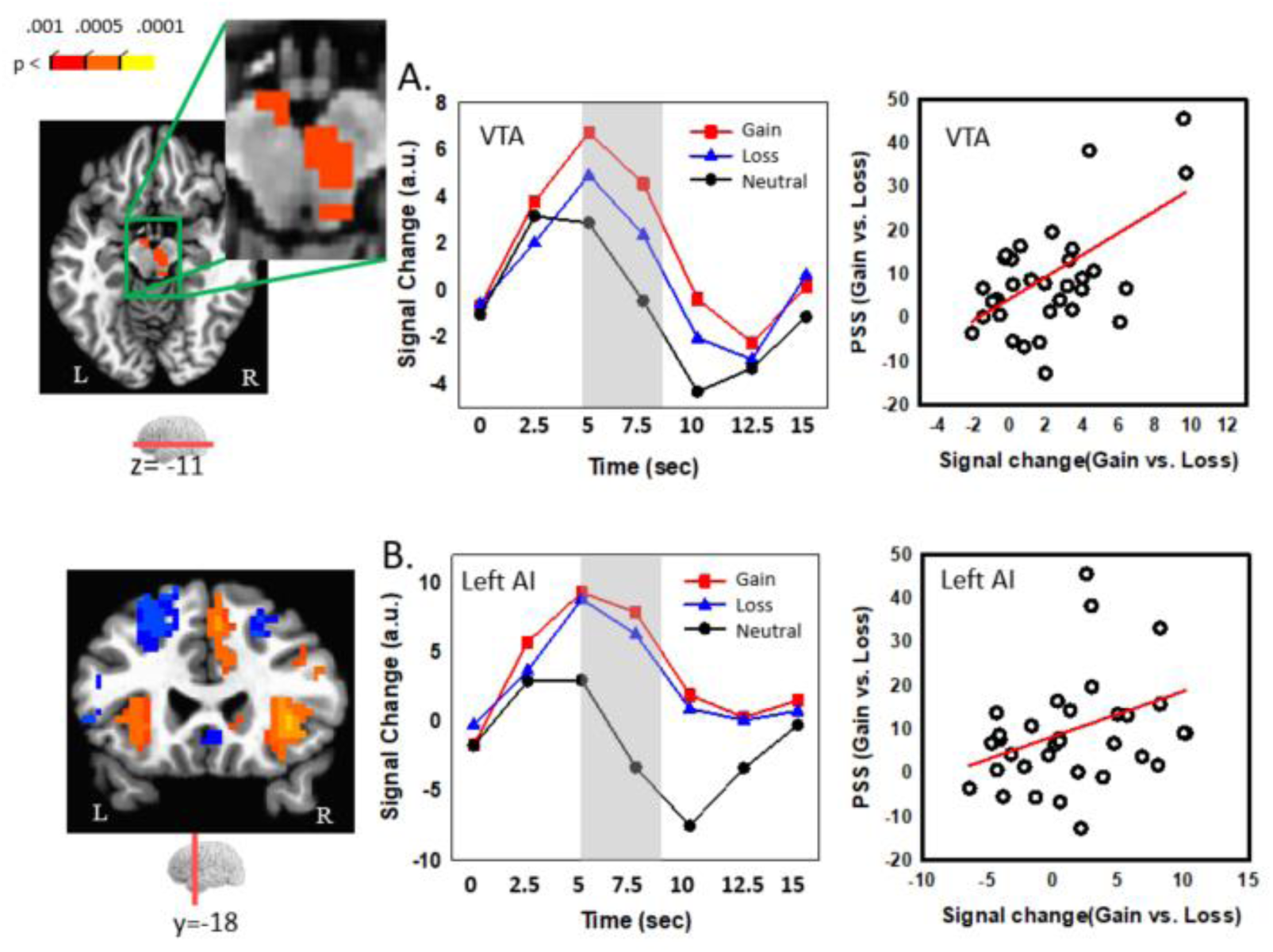
Individual difference in mesolimbic response predicts bias in temporal perception of gain and loss-associated cues. Average deconvolved responses (left panel), mean response indices as a function of time (middle panel) during monetary incentive task, and association with temporal order judgment asymmetry, as indexed by point of subjective simultaneity, PSS (right panel). Greater PSS reflects greater distortion in temporal perception. Top panel, ventral tegmental area (VTA); Bottom panel, left anterior insula (AI).

We find that Individual differences in motivational orientation and reward system response both contribute to temporal distortions of gain relative to lose cues. To better understand the relationship between personal values and reward response, we conduct two mediation analyses. While individual differences in VTA response do not correlate with regulatory focus (Pearson’s r =.22, p=.22; robust R^2^ = .04, p=.26), the relation between the anterior insula and temporal bias, PSS (gain vs. lose), is explained by both an individual’s VTA tuning (gain vs. loss) and regulatory focus (promotion vs. prevention) (**Fig. 6AB**). Individual differences in motivational orientation and VTA response contribute to anterior insular salience(Anderson et al., 2011), biasing the subjective temporal priority of cues signaling gain over loss opportunities.

**Figure 6.**
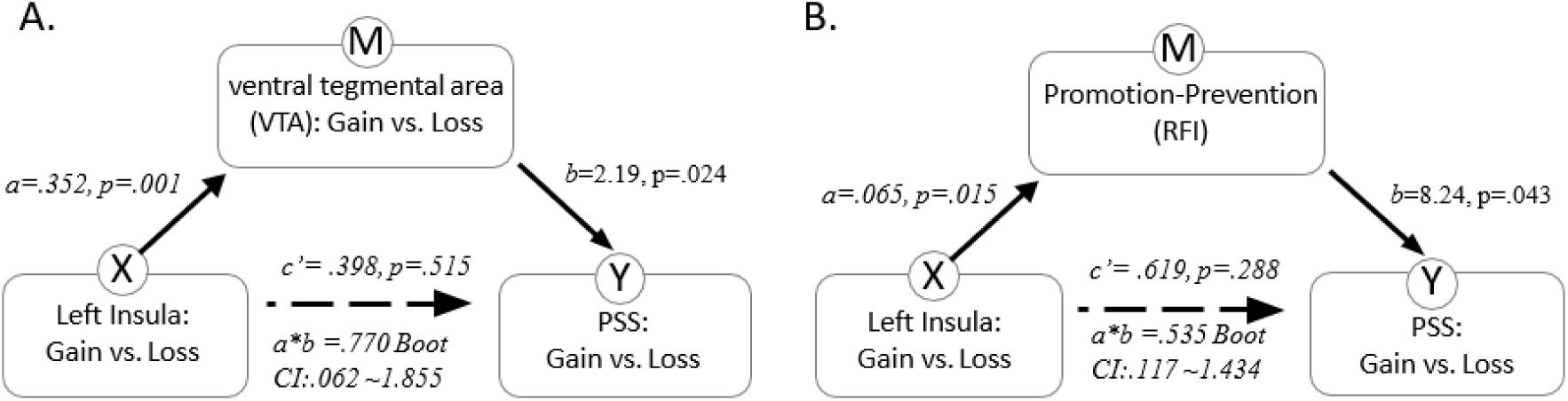
Motivational and mesolimbic mediation of distortion in gain and loss (loss avoided) temporal perception. The dotted line indicates the adjusted path coefficient c’ between Insula (X) and PSS (Y) is reduced to nonsignificance after the mediator (A, VTA signal; B, regulatory focus, RFI) is considered. The letters *a, b, a*b,* and *c’* refer to estimated path coefficients. Boot CI is the bootstrapped confidence interval.

## Discussion

Prospect theory has suggested that decisions reflect a bias toward avoidance of loss compared to equivalent gains(25, 64, 65). A common interpretation of loss aversion is that losses carry greater subjective importance than gains, such as losses producing more attention and attracting more emotion (13, 36). Previous research is mostly carried out in high-level decision-making domains and is largely driven by the unequal probability between gain and loss. Unlike previous work, in the present studies, gain and loss are equally reinforced. We investigate whether a loss-associated stimulus is given the same priority related to a gain-associated stimulus in perception, which allows us to test the theoretical models and understand the fundamental question of where the loss aversion comes from.

In Study 1, we test the ***attention-based hypothesis*** of loss aversion. Among n=148 participants, there is a general attentional bias toward gain over loss-related cues, which results in the distorted temporal perception of color cues associated with these opportunities. When faced with equal potential gains or averted losses, events predicting gains win the head-to-head race for being perceived first. Loss does not produce more attention, which sort of eliminates the explanation for loss-averse behavior that losses draw attention more than gains. In Study 2, we scan participants and test the ***contrast-based hypothesis*** of loss aversion. We show that brain regions respond similarly to gains and losses, including the VTA, NAcc, anterior insula, and mPFC. Previous studies show that losses give rise to more activation in emotional regions, such as the amygdala and anterior insula(30–34). In the present study, amygdala activation to losses is not observed, and for most participants, gains produce more activities in the emotion-related areas, such as VTA and anterior insula, eliminating that losses produce more emotion. Across two studies, we show systematic individual differences in motivational orientation, reward system response, and inattention to lose than gain (loss posteriority).

Loss aversion suggests that a loss has more influence on choices than a gain of the same magnitude. It is certainly the most significant contribution of psychology to behavioral economics(66). However, in a series of experiments, losses generate neither more attention nor more emotion than gains. The absence of the asymmetry of loss vs. gain in directing attention leads to **a basic question of how then risk aversion arises**. It must be something like a higher level consideration of what it might mean to lose something that people already have(67, 68). This is implicated in the current literature. **First**, loss aversion exists in the higher level of comparisons(37–42). Specifically, loss aversion appears only when gain-loss comparisons are encouraged(69), and it is more pronounced in loss contemplation(70), affective simulations, or forecasting(71, 72). It has shown that risk preferences are shaped by couture and malleable in response to new environments(73). **Second,** loss aversion appears when people expect the pain of losing what they gain to exceed the pleasure of gaining. The *endowment effect,* perhaps, is the most prominent illustration of the consequences of loss aversion. In 1991, Kahneman and colleagues noted that “the main effect of endowment is not to enhance the appeal of the good one owns, only the pain of giving it up”(78, p.197). This higher-order processing could occur consciously and or unconsciously(74, 75), reflecting anchor points in self-regulation(47, 76). Naturally, the differences in considering of losses vs. gains relate to and guide loss aversion behavior across subjects.

The present research also builds on the work by associating loss aversion to self-regulatory mechanisms. Theories have suggested that the strategy one adopts in decisions may vary in its association with approach vs. avoidance-related behaviors(22, 23), and individuals possess trait-like dispositions, called regulatory foci(48). RF theory posits two distinct motivation systems and thus goes beyond the original reference point’s hypothesis in prospect theory(25). Regulatory focus indexes behavioral-regulation strategies formed by the nurturing environment, such as attachment, parenting styles, and patriarchal culture. As seen in Fig. 3C, prevention-focused individuals tend to exhibit the most distorted temporal perceptions to gain, loss aversion; and by contrast, promotion-focused ones exhibit the most distorted temporal perceptions to loss, loss posteriority. Thus individual differences in loss-gain contrast vary along the self-regulatory dimension of goal commitment. Consistently, in the brain, we identified voxels within the two functional ROIs (i.e., VTA and insula) in which changes in BOLD signal during loss relative to gain are correlated with participants’ attentional bias of gain over loss. The greater responses in VTA and insula to loss relative to gain during monetary reinforcement, the greater disparity in their temporal perception of loss vs. gain. ***Specifically***, the path modeling (mediating analysis) confirms our mechanisms hypothesis that the ability of insula to predict distorted perception of gain vs. loss is mediated through the impact of VTA and RF orientation. In sum, individual differences in motivational orientation and reward system response both contribute to the contrast of gain vs. loss in perception.

Ultimately, loss aversion relates to higher level consideration and can be seen as a stable individual trait and or as a response to particular circumstances(73). Further, intentional cognitive regulation, e.g., thinking like a trader, modulates loss aversion(42), and once the pre-attitude is removed, loss aversion is reduced or eliminated(77). These all seem to suggest that individual intentions are moderators of loss aversion in general(78). In fact, people’s intention is the determinant of the reference points, and that reference points reflect expectations is part of the original prospect theory(47). For instance, Loss aversion is observed in those with a predominant prevention focus but not in individuals with a promotion focus(29, 79, 80), and negative outcomes are more motivating in a prevention focus than in a promotion focus, whereas the reverse is true for positive outcomes(29, 80). A recent study suggests that the pre-valuation bias, a kind of regulatory processing, determines loss aversion(81).

We show that most individuals demonstrate asymmetric attention toward cues for gains over losses in directing attention, and this is supported, in part, by neural differences originating from midbrain VTA responses and their influence over the anterior insula - most people show a greater emotional response to gains, not losses in critical emotion and motivation regions. The loss posteriority in perception may be a basic and far-reaching psychology principle. The VTA differentially shapes attention and awareness to gains and losses, and individual differences in behavioral and neural loss posteriority observed in the present study are related to naturally occurring differences in the mesocorticolimbic dopamine systems. Our findings would have implications for a wide range of situations. For instance, the weight difference of gain over loss in perception should be considered in marketing, brand development, and design. Without such understanding, well-intended policies may be ineffective. As a general-purpose mechanism of attentional capture, the increased sensitivity to gains of some participants may have contributed to their elevated attentional priority to gain in perception(48, 54). Specifically, the maladaptive attentional biases found in drug addiction, obesity, and anxiety(82, 83) may reflect the disordered influence of the time inconsistency and asymmetry of gain and loss in perception. Note that the VTA and insula are common core regions for psychiatric illness and contribute to mental states(84). RFI on affective disorders has received less attention, while shifts of the regulatory focus and or reframing a given set of outcomes as relative gain over relevant losses could lead to systematic reversals of gain-loss competition and risk preferences over that set.

These data suggest a neurobiological link between human loss aversion behavior and regulatory focus in perception. While attention and emotion are key components in decision making, our results show that loss aversion does not rest on attention or emotion, and losses may be less accessible to the extent if it is not perceived or framed as gains for most people.

## Methods

For additional information, please see *SI Text*.

### Participants

There are 148 participants involved in Study 1 (87 females, mean age =22) and 34 participants involved in Study 2 (17 females, mean age =22). All are from an ongoing data collection project^1^: The experiments are similar, and each consists of two intermixed trial types: *Side judgment* and *temporal order judgment* (TOJ) trials. All are undergraduate and graduate students from Cornell University, Ithaca, and provide informed consent, as approved by the Institutional Review Board of Cornell University.

**Fig.1** illustrates the core tasks and the examples of the stimuli. The stimuli consisted of luminance-matched colored circles. Computer screen color is manipulated using HSL scheme: blue (160,240,120; corresponding RGB: 0, 0,255), red (0,240,120; corresponding RGB: 255, 0, 0), and yellow (40,240,120; corresponding RGB, 255, 255, 0). The background was set to grey (HSL: 170, 0, 84; RGB: 84, 84, 84). (*SI Appendix, Supplementary Methods*).

For the value reinforcement task, each trial starts with a fixation display for 1000 ms and is followed by a circle with a left or right gap, in red, yellow, or blue for 800 ms. Participants are required to indicate “left gap” or “right gap” by pressing a corresponding button as quickly and as accurately as possible. Before the start of the experiment, participants are informed that with one of the color categories, there would be an opportunity to gain or lose (prevent the loss) money based on performance (100% contingency, also informed to participants). Feedback followed each trial indicating money gained, missed, saved or lost. During the temporal order judgment (TOJ) trials, participants are instructed to report “which side came first?”, by pressing a corresponding key during the TOJ task. Unlike the value reinforcement task, a speeded response is not emphasized. A stimulus onset asynchrony (SOA) separates the first and second circles by 8,18,38,68, and 98 ms. Circles are presented until a response (max, 1.5s).

The experiment includes 5 runs, and each run consisted of 60 value reinforcement trials (20 gain, 20 loss, and 20 neutral) and 60 TOJ trials. Each run for the TOJ task utilized a 5 SOAs x 6 (value pairs:

Gain-Neutral, Neutral-Gain, Loss-Neutral, Neutral-Loss, Gain-Loss, Loss-Gain; note the first-mentioned of each pair appears first) x 2 (the first appearing stimulus is presented on the left vs. right side) design. Color value pairing is counterbalanced across participants. The schedule of stimulus presentation and data collection are controlled by Presentation software (Neurobehavioral Systems, Albany, CA).

### Behavioral data analysis

We fit a Gaussian function to each individual’s data across onset asynchronies from the ***TOJ task*** (49). PSS indicates how much prior in time a stimulus would need to be shown to result in a subjective perception of simultaneity(85, 86), with the null hypothesis being 0 ms, representing no temporal distortion.

### fMRI study

MR data are collected using a multi-echo 3 Tesla GE Discovery MR750 (Independent receiver channel: 32). For each functional run, BOLD EPI volumes are acquired with a TR of 2600 and TEs of 13.7, 30, and 47 ms. fMRI data are analyzed using tools from the AFNI software package(87). Multi-echo fMRI uses ME-ICA to create a weighted combination of three echoes to form a single time series, which we have found enhances discrimination of small nuclei in high susceptibility regions(88). Finally, regression analysis is performed on these data, and the alpha level for voxelwise statistical analysis is determined by simulations using the 3dClustSim program of the AFNI toolkit(89). Considering the report of increased false-positive rates linked to the assumption of Gaussian spatial autocorrelation in fMRI data(90), we use the –acf (i.e., autocorrelation function) option added to the 3dFWHMx and 3dClustSim tools. We examine the neural bases of individual differences in TOJs for gain and lose cues by linear regression of a set of regions of interest (ROIs) derived from the independent value reinforcement task. Reward ROIs are defined based on the main effect of reward at a cluster-extent threshold of 32 voxels (p<.005, uncorrected), independent from the TOJ performance (k=32 contiguous voxels for a corrected threshold of p<.05).

## Authors’ notes and Acknowledgements

Dr. Michael Posner contributes significantly to this project. We extend our sincere gratitude for his invaluable input and guidance throughout the development of this paper. His work encompasses project supervision, conceptualization, writing, review, and editing.

## Footnotes

1 Our paper studies the source of loss aversion behavior and its variability, specifically, the neurobiological link between human loss aversion behavior and regulatory focus in perception. Behavior reports have used a small sample from the current project for other questions.

