## Supplemental file for "Attentional priority and limbic activity favor gains over losses"

**Correspondence:**

### Supporting Information

#### Study 1: Loss and attention

##### *Methods*

***Participants.*** There are six experiments with n=148 participants involved in this study (87 females, mean ages =22). *The data are from an ongoing data collection project.* We select a similar testing procedure for such experiments: The conditions include gain vs. loss value reinforcement followed by a temporal order judgment. All are undergraduate and graduate students from Cornell University, Ithaca, and provided informed consent, as approved by the Institutional Review Board of Cornell University. Subjects are free from psychiatric or neurological disease or related history, as indicated via self-report. All report normal or corrected-to-normal acuity and color vision, and all are naive to the purpose of the experiment. All are paid immediately after the experiment.

***Stimuli and behavioral paradigm*** Fig. 1 illustrates the core tasks and the examples of the stimuli. The stimuli consist of luminance-matched blue, red, and yellow circles (size: 240 x 240 $px^2$, 5.5 $lm/ft^2$). Extech Foot Candle light meter is used to measure color illuminance within the field of view. Computer screen color is manipulated using HSL scheme: blue (160,240,120; corresponding to RGB: 0, 0,255), red (0,240,120; corresponding to RGB: 255, 0, 0), and yellow (40,240,120; corresponding to RGB: 255, 255, 0). The background is set to grey (HSL: 170, 0, 84; RGB: 84, 84, 84).

During the reinforcement trials, participants are instructed to perform a side discrimination task. Each trial starts with a fixation display for 1000 ms and is followed by a color circle (left or right gap, in red, yellow, or blue) for 800 ms. There is a jittered 1 – 5 second inter-stimulus interval (ISI, mean: 2 sec) containing a blank screen that appeared after the target display. Participants are required to “left gap” or “right gap” by pressing a corresponding button as quickly and as accurately as possible. Before the start of the experiment, participants are informed that with one of the color categories, there would be an opportunity to earn or prevent the loss of money based on performance (100% contingency, also informed to participants). On each earn trial, participants win $0.30 if they are both accurate and fast. They win $0.00 during error or slow trials. During loss trials, participants avoid loss if they are both accurate and fast; otherwise, they lose $0.30 for either slow or error trials. During neutral trials, participants do not gain or lose money regardless of their performance. At the end of each value reinforcement trial, a visual feedback display (duration: 1 sec) provides the accumulated “earn” and “save” bars; concurrently, a sound is played acknowledging participants' gain (beep sound 1) or loss (beep sound 2) of money. A different sound (sound 3) is played if participants provide an incorrect response. The RT threshold for a “fast” response is set between 350-650 ms.

During the TOJ trials, each trial starts with an initial fixation display for 1000 ms. Subsequently, a colored circle (no gap, either in red, blue, or yellow) is presented either on the left or on the right side of the fixation; this is, followed by a second circle of a different color, which appears on the opposite side of the first circle after a randomly determined stimulus onset asynchrony (SOA: 8, 18, 38, 68, and 98 ms). During TOJ trials, participants are instructed to report either which color or side stimulus appears first by pressing a corresponding key during the TOJ task. Stimuli remain on the screen until response. Color-gain and lose (loss avoidance) associations are counterbalanced across participants. Unlike the value reinforcement task, the speeded response is not emphasized for the TOJ task. Finally, a jittered 1 – 5 second jittered inter-trial interval (ITI, mean:2 sec) containing a blank screen terminates the trial. Participants are informed that all TOJ trials are neutral (i.e., providing no chance of gain or loss). While TOJ stimuli are closed circles, during the reinforcement trials, the stimuli have a gap on either the left or right side. This allows us to ensure color is affected by gain, loss, and neutral coding (color–value association) from prior reinforcement training. The experiment includes 5 runs, and each run consists of 60 value reinforcement trials (20 earning, 20 saving, and 20 neutral) and 60 TOJ trials. The visual perception task begins with a value reinforcement trial, and then, the reinforcement trials pseudo-randomly alternate with TOJ trials. Each run for the TOJ task utilizes a 5 (SOA separating first and second stimulus: 8,18,38,68 and 98 ms) x 6 (value pairs: Earn-Neutral, Neutral-Earn, Loss-Neutral, Neutral-Loss, Earn-Loss, Loss- Earn; note the first mentioned of each pair appears first) x 2 (the first appearing stimulus is presented on the left vs. right side) design.

The schedule of stimulus presentation and data collection are controlled by Presentation software (Neurobehavioral Systems, Albany, CA). Each participant is given a practice run of 30 trials not analyzed. Among this practice run, one-half are discrimination task trials, and the other half are TOJ task trials.

**Regulatory focus Questionnaire** Participants' focus is measured with an 11- item Regulatory Focus Questionnaire (RFQ(1)). A promotion focus is associated with seeking advancement and accomplishment, while the prevention focus corresponds to the concerns of safety and responsibility. The promotion subscale contains six items (e.g., “Do you often do well at different things that you try?”), while the prevention scale contains five items (e.g., “How often did you obey rules and regulations that were established by your parents?”). Responses are made on a 5-point rating scale (1 = never or seldom, 5 = very often), and the prevention subscale total is subtracted from the promotion subscale total to create the regulatory focus index (RFI)(2).

***Data analysis and results***

Performance in the value reinforcement task is measured by participants' reaction times (RTs), error rates, and proportions of gain receipt and successful loss avoidance. For each participant, mean RTs for the correct response for each condition are calculated. RT and error rate data are submitted to a one-way repeated ANOVA.

To assess the TOJ task, we compute the mean proportion of trials in which the target is judged as "appearing first" in two conditions: 1) when it physically appears first (denoted by SOA: 8, 18, 38, 68, and 98 ms); 2) when it physically appears second (denoted by SOA: -8,- 18, -38, -68, and -98 ms). We assess the magnitude of temporal distortion for "Gain vs. Neutral ", "Loss vs. Neutral", and "Gain vs. Loss", by calculating each participant's point of subjective simultaneity (PSS), which indicates the time interval needed by participants to perceive the two stimuli as arriving simultaneously(3–6). PSS is calculated using Gaussian fit method (the Gaussian function from Matlab, Mathworks, MA, USA). For Experiment 1 (n=36), prior entry is observed for gain and loss, also reflected on participants on average perceive gain and loss stimuli at arriving 12.9 and 6.2 ms (PSS) prior to neutral stimuli, respectively, for gain vs. neutral, $t(35) = 5.51$, $p<.001$, Cohen's $d=.92$; and for loss vs. neutral, $t(35) = 2.99$, $p=.005$, Cohen's $d=.50$, and the effect for gain is larger than that for loss, $t(35) = 2.91$, $p=.006$, Cohen's $d=.48$.

#### Study 2: Loss and emotion

##### *Methods*

***Participants.*** Thirty-four Participants participate in the experiment (17 males, mean age =21, all right-handed) and provide informed consent, as approved by the Institutional Review Board of Cornell University, Ithaca, NY. One participant is excluded from further analysis due to his slow responses (z=3.37) in the TOJ task. Subjects are free from psychiatric or neurological disease or related history, as indicated via self-report. All report normal or corrected-to-normal acuity and color vision, and all are naive to the purpose of the experiment. All are paid immediately after the experiment. All are undergraduate and graduate students from Cornell University, Ithaca, and have self-reported normal or corrected-to-normal vision and intact color vision.

**Self-reported affect.** To examine potential color preferences and their modulation following value reinforcement, we assess subjective valence and arousal responses to each of the color stimuli using 1-7 Likert scales (not at all" to "extremely", 1 to 7) before and after the experiment.

***Stimuli and behavioral paradigm*****.** Methods are similar as experiments reported above. We set a strict RT threshold (400 ms) for successful gain and loss avoidance, avoiding a ceiling effect for value reinforcement.

*MRI Data Acquisition*. MR data are collected using a multi-echo 3 Tesla GE Discovery MR750 (Independent receiver channel: 32). The scanning session begins with a high-resolution T2-FLAIR anatomical scan (TR = 1200 ms, TE = 9.5 ms, TI =271.2 ms, 0.3 mm isotropic voxels). Then for each functional run, BOLD EPI volumes are acquired with a TR of 2600 and TEs of 13.7, 30, and 47 ms. Each volume consists of 42 oblique slices with a thickness of 3 mm and an in-plane resolution of 3 X 3 mm (21.6 mm field of view). We employ Multi-Echo ICA(7) to reconstruct brain volumes differential echo weighting to better identify a regional BOLD response.

#### General ***Data Analysis***

Examining reinforcement performance, on average, participants have an equal proportion of gain (56%, \$16.8 gained) and loss (57%, \$17.2 loss avoided), t(33)=1.51, p=.140, Cohen's d=.22, at the end of the experiment.

There is a trend for reinforcement on response time, $F(2,55)=3.00$, $p=.067$, $\eta^2=.08$, reflecting that responses on gain and loss trials are faster than neutral. The ANOVA for error rate shows a significant main effect, $F(2,50)=13.52$, $P<.001$, $\eta^2=.29$, revealing greater accuracy on neutral relative to gain and loss trials, $t(33)=5.12$, $p<.001$, Cohen's d=.95 (SI **Fig.2 A**), consistent with the speed-accuracy tradeoff. Examining participants' temporal order judgments, we also analyze accuracy, submitting hit rates to ANOVA [temporal order (Gain-first, Loss-first) x SOA (8, 18, 38, 68 and 98 ms)]. A main effect of SOA, $F(3,95)=104.34$, $P<.001$, $\eta^2=.76$, reveals that participants are much more accurate when the temporal difference between colored circles is greater, and on average for gain-first relative to loss-first colors, $F(1,33)=16.36$, $P<.001$, $\eta^2=.33$. A significant two-way interaction, $F(3, 95)=10.75$, $P<.001$, $\eta^2=.25$, reveals lower hit rates for loss-first relative to gain-first colors is most pronounced at short SOAs, with accuracy at the shortest SOA on loss-first trials not differing from chance (SI **Fig.2 B**). Loss cues are thus relatively unattended compared to gain cues, resulting in poor temporal judgment performance.

To examine relative temporal perceptions, we fit a Gaussian function to each individual's temporal order data across onset asynchronies (3, 8) to estimate the point of subjective simultaneity (PSS) for gain and loss colors. Participants perceive both gain and loss colors as arriving prior to neutral, respectively: $PSS_{Gain-Neutral}$, $t(33)=4.09$, $p<.001$, Cohen's d=.70; $PSS_{Loss-Neutral}$, $t(33)=3.07$, $p=.004$, Cohen's d=.53(**SI, Fig. 3A**). When gain and loss colors compete head-to-head, gain stimuli are judged to appear before the concurrently presented loss stimuli, $t(33)=3.90$, $p<.001$, , Cohen's d=.67, consistent with the greater acquired salience of cues predicting gaining. Once again, there are substantial individual differences in the expression of this temporal distortion. In this subsample, regulatory focus orientation is associated with temporal perceptions, Pearson's r= .48, p=.007; robust $R^2$ =.23, p =.009, with increased promotion relative prevention, focus associated with reduced loss salience (**SI, Fig. 3B**).

In the present studies, significance level for main statistical analyses is set at $p < 0.05$. The Greenhouse–Geisser correction is used when the sphericity assumption is not met in ANOVAs(9). Planned comparisons are then performed to test the value reinforcement effect between gain and loss. Holm-Bonferroni correction was applied to the alpha criterion for multiple comparisons when determining significance. As elsewhere(10, 11), we run an across-subject robust regression analysis (the *robustfit* function from Matlab, Mathworks, Natick, MA, USA), given that standard Pearson correlation is sensitive to even a few influential data points(12); results from robust regression were reported in terms of $R^2$. When necessary, we use Matlab function *rcoplot* to examine potentially influential data points (confidence intervals 95%). Following (13), effect sizes are reported as $\eta^2$ (small, 0.01; medium, 0.06; large, 0.14) for ANOVAs, and as d (small = 0.20; medium = 0.50; large = 0.80) for planned comparison t-tests.

##### fMRI Data Analysis

fMRI data are analyzed using tools from the AFNI software package(14). For the ME-ICA analysis pathway, the following preprocessing is performed on unprocessed time series data(15). The first 3 volumes of each functional run are discarded to account for equilibration effects. Slice time correction is applied (3dTShift). Motion correction parameters are estimated for each time point by aligning the middle TE (30 ms) images to the corresponding first-time point image using a rigid-body (6 parameters) alignment procedure (3dvolreg). The functional to structural co-registration parameters are estimated by registering the skull-stripped middle TE image from the first time point to the skull-stripped anatomical image using an affine (12 parameters) alignment procedure with the local Pearson correlation cost-function (3dSkullStip, 3dAllineate)(16). Motion correction and anatomical co-registration parameters are then applied in one step (3dAllineate). A brain mask is computed from the mean image of the shortest TE (13.7 ms) time series (3dskullstrip) and applied to all images. Each image is spatially smoothed with a 5 mm FWHM Gaussian kernel within the functional mask (3dBlurInMask). As typical for multi-echo fMRI data, the three echoes are combined to form a single time series. Finally, regression analysis is performed on these data (see below).

Whole-brain voxelwise random-effects analyses are restricted to gray-matter voxels based on the FSL automated segmentation tool FAST (FMRIB's Automated Segmentation Tool; http://www.fmrib.ox.ac.uk/fsl/). We consider three main event types in the design matrix: gain, save and neutral trials. Constant, linear, and quadratic terms are included for each run separately (as covariates of no interest) to model baseline and drifts of the MR signal. To account for the signal variance related to head motion, six estimated motion parameters are included as nuisance regressors in the model. No assumptions are made about the shape of the hemodynamic response function. Responses are estimated starting from picture onset to 15 sec using cubic spline basis functions. As an index of response strength, we average the estimated responses at 5 and 7.5 s after stimulus onset (as determined via spline-based estimates) for all main event types.

The first goal of the voxelwise analysis in the discrimination task is to define regions of interest (reward versus neutral). Whole-brain voxelwise random-effects analyses are conducted using response estimation from individual-level analyses (restricted to gray matter voxels) in AFNI. To probe the effects of the regressors of interest, we run separate one-sample t tests again zero using the AFNI's 3dttest ++ program. The resulting statistical maps were thresholded at $p<.005$, and we corrected for multiple comparisons using 3dClustSim program. The alpha level for voxelwise statistical analysis is determined by simulations using this program (restricted to gray matter voxels). For these simulations, the smoothness of the data is estimated using 3dFWHMx program (restricted to gray matter voxels) based on the residual time series from the individual-level voxelwise analysis. Considering the report of increased false-positive rates linked to the assumption of Gaussian spatial autocorrelation in fMRI data(17), we use the –acf (i.e., autocorrelation function) option added to the 3dFWHMx and 3dClustSim tools, which models spatial fMRI noise as a mixture of Gaussian plus mono-exponential distributions. This improvement is shown to control false-positive rates around the desired alpha level, especially with relatively stringent voxel-level uncorrected p values such as .005(18).

Each activation cluster of the main effects themselves is defined as an individual ROI. We adopt the selection criterion of main effects to determine ROIs, because it is statistically independent of the central goal of our analysis, that is, to probe distinct effects of gain and save on temporal order judgments(19, 20). A representative time series for each ROI is then created by averaging the time series of all the gray matter voxels inside the ROI (5-mm radius). Then, for each ROI, as in the whole-brain voxelwise analysis, deconvolution analysis is run on the representative time series data to estimate the hemodynamic response function of two main regressors of interest. Then, for each condition, we take the estimated mean response at 5 and 7.5 sec time points following stimulus onset as the index of response strength.

#### Mediation Analysis

To test the models of reward response and motivational orientation relate to temporal perceptions displayed in Fig. 4, we perform mediation analysis(21, 22). A standard mediation model involves evaluating these four components: 1) total effect *C* (initial variable → outcome, which equals to the direct effect *c′* plus the indirect effect *a*b*); 2) indirect path *a* (initial variable → intervening variable, mediator; 3) indirect path *b* (intervening variable → outcome, after controlling for the initial variable; and 4) direct effect *c′* (initial variable → outcome, after controlling for the intervening variable). The final mediation effect is tested by assessing the significance of the product of paths a and b by using bootstrap tests (23, 24).

**Supplementary Figure 1**

**Individual differences in money gained versus saved in the value reinforcement task.** Panel A: Frequency distribution of temporal judgments for money gained vs. saved colors (i.e., \$(Gained) - \$ (Saved); Panel B: Bayesian T-Test: Prior and posterior information.

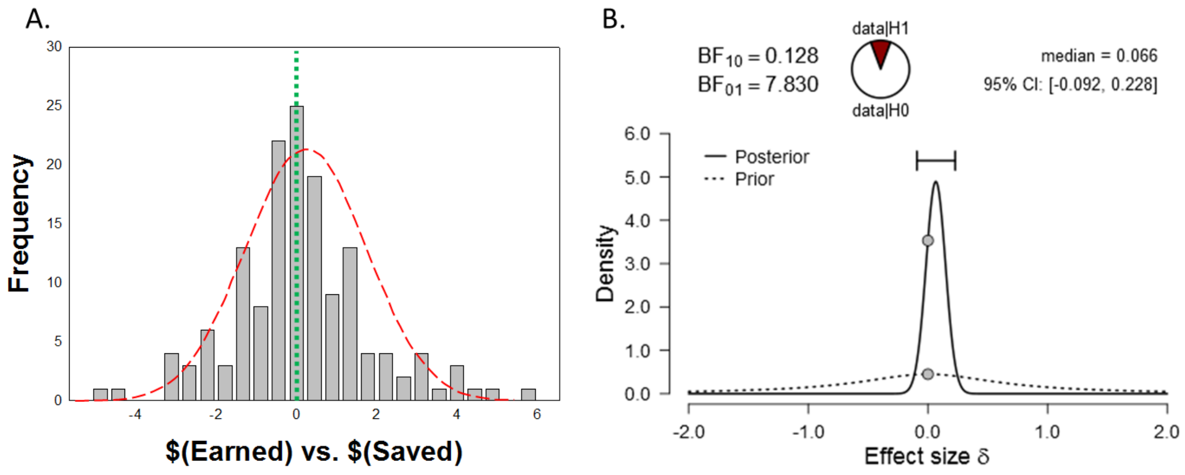

**Supplementary Figure 2**

**Behavioral data for the fMRI study.** Panel A: RTs and error rates for the value reinforcement task; Panel B: Accuracy for the TOJ task.

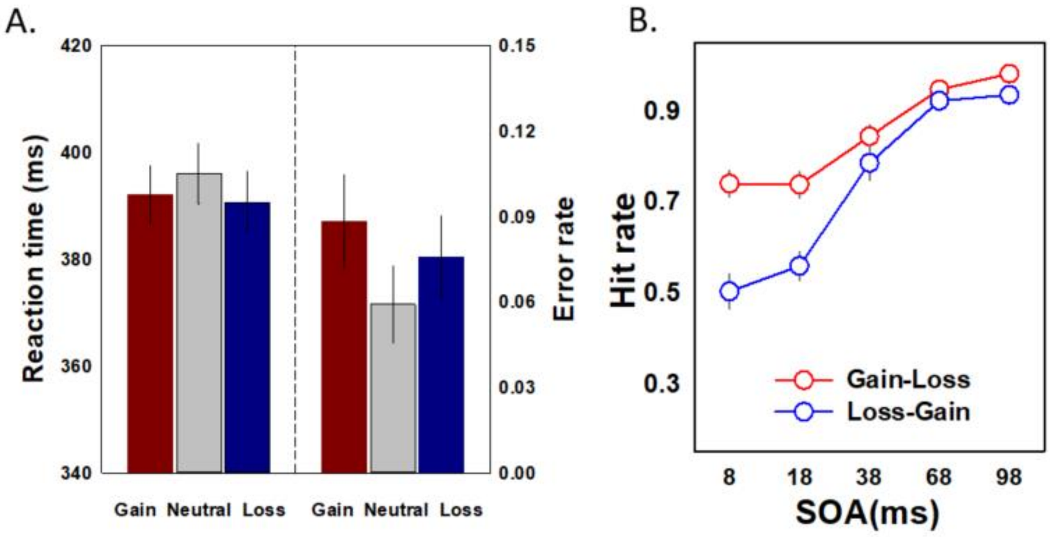

**Supplementary Figure 3**

**Temporal order judgments for gaining and saving cues and their relation to motivational orientation.** Panel A: Point of perceived simultaneity (PSS) sizes were estimated using Gaussian function for gain vs. neutral, gain vs. loss, and loss vs. neutral trials. Panel B: Individual differences in regulatory focus index (RFI, promotion minus prevention) were related to the magnitude of the gain-loss asymmetry in temporal order judgment, i.e., PSS (Gain vs. Loss) scores. Dot lines show the confidence intervals 95%.

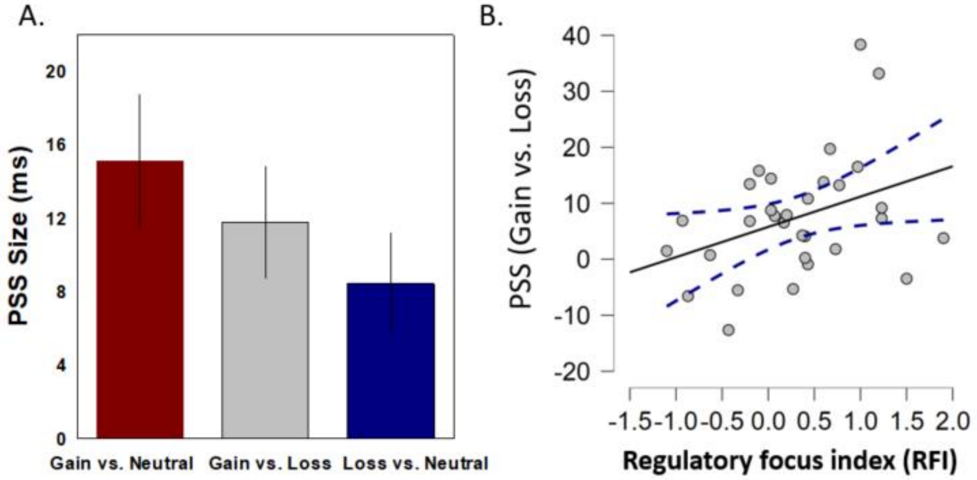

**Supplementary Figure 4**

**Changes in affective response to colors following gain and lose value reinforcement.** Panel A: Association between an individuals' signal change in insula (Gain vs. Lose) and self-reported ratings of valence to gain and loss colors. Panel B: Association between an individuals' signal change in the insula (Gain vs. Lose) and VTA (Gain vs. Lose).

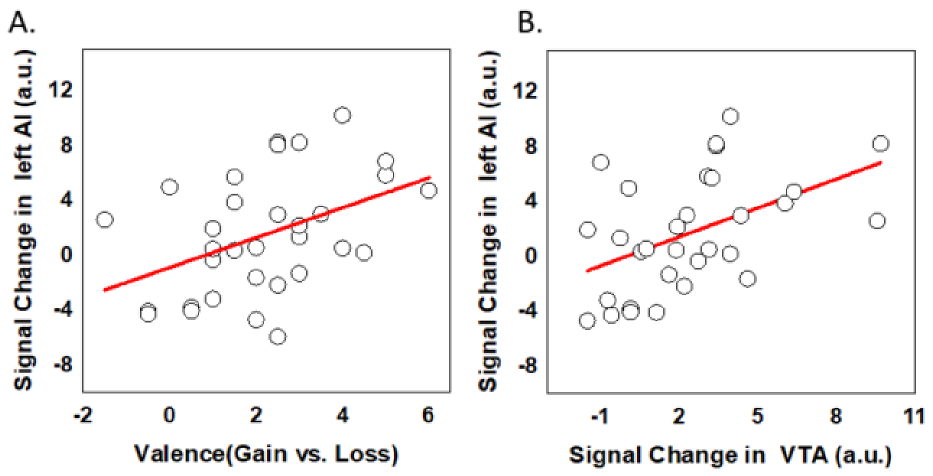

**Supplementary Figure 5**

Results of ROI (right anterior insula) and correlation analysis. Average deconvolved responses (left panel), mean response indices as a function of time (middle panel), and association with temporal asymmetry in the point of subjective simultaneity, PSS (right panel). See text for details.

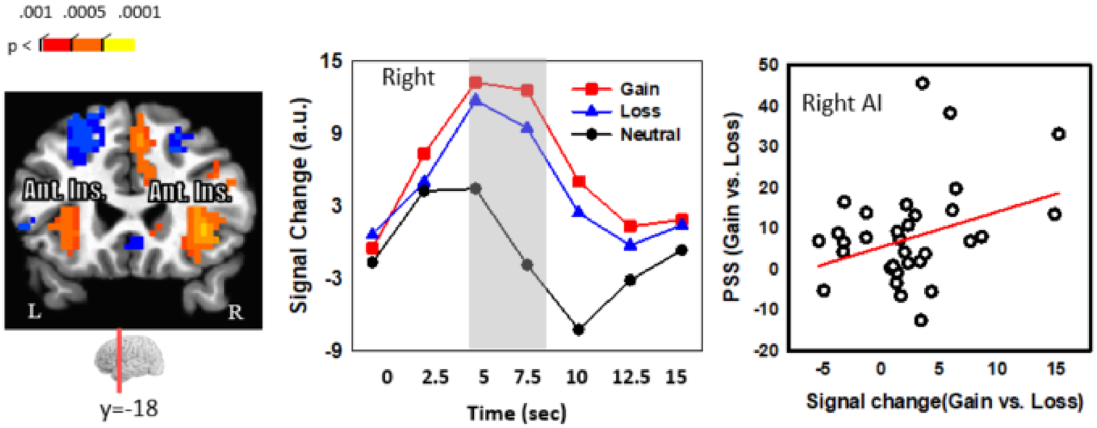
